# Cycling Physicochemical Gradients as ‘Evolutionary Drivers’: From Complex Matter to Complex Living States

**DOI:** 10.1101/000786

**Authors:** Jan Spitzer

## Abstract

**Highlights:**

- Biological complexity cannot be reduced to chemistry and physics
- Complex living states are: multicomponent, multiphase, ‘crowded’, and *re-emergent*
- Living states arise naturally *only* by the action of cycling physicochemical gradients
- Bacterial cells can be modeled as viscoelastic capacitors with sol-gel transitions
- Evolving living states can be investigated via ‘biotic soup’ experimentation
- Darwinian evolution arises from the process errors of the cell cycle
- Synthetic biology heralds the transition from unintentional Darwinian evolution to intentional anthropic evolution

**Abstract:** Within the overlap of physics, chemistry and biology, complex matter becomes ‘more deeply’ understood when high level mathematics converts regularities of experimental data into scientific laws, theories, and models (Krakauer et al., 2011. *The challenges and scope of theoretical biology*. J. Theoret. Biol. 276: 269–276). The simplest kinds of complex biological matter are bacterial cells; they appear complex–from a physicochemical standpoint–because they are multicomponent, multiphase, biomacromolecularly crowded, and re-emergent; the property of re-emergence differentiates biological matter from complex chemical and physical matter.

Bacterial cells cannot self-reassemble spontaneously from their biomolecules and biomacromolecules (via non-covalent molecular forces) without the action of external ‘drivers’; on Earth, such drivers have been diurnal (cycling) physicochemical gradients, *i.e*. temperature, water activity, etc. brought about by solar radiation striking the Earth’s rotating surface. About 3.5 billion years ago, these cycling gradients drove complex chemical ‘prebiotic soups’ toward progenotic living states from which extant bacteria evolved (Spitzer and Poolman, 2009; *The role of biomacromolecular crowding, ionic strength and physicochemical gradients in the complexities of life’s emergence*. Microbiol. Mol. Biol. Revs. 73:371–388). Thus there is historical non-equilibrium continuity between complex ‘dead’ chemical matter and complex living states of bacterial cells. This historical continuity becomes accessible to present-day experimentation, when cycling physicochemical gradients act on ‘dead’ biomacromolecules obtained from (suitably) killed bacterial populations – on a ‘biotic soup’ of chemicals (Harold, 2005, *Molecules into cells: specifying spatial architecture*. Microbiol. Mol. Biol. Rev. 69:544–564). The making of biotic soups and recovering living states from them is briefly discussed in terms of novel concepts and experimental possibilities.

In principle, emergent living states contingently arise and evolve when cycling physicochemical gradients continuously act on complex chemical mass; once living states become dynamically stabilized, the inevitable process errors of ‘primitive’ cell cycles become the roots of Darwinian evolution.

---

> (i) In physical science a first essential step in the direction of learning any subject is to find principles of numerical reckoning and practicable methods for measuring some quality connected with it. I often say that when you can measure what you are speaking about, and express it in numbers, you know something about it; but when you cannot measure it, when you cannot express it in numbers, your knowledge is of a meager and unsatisfactory kind; it may be the beginning of knowledge, but you have scarcely in your thoughts advanced to the stage of science, whatever the matter may be.
>
> — Lord Kelvin (1883)

> (ii)…we have learned to appreciate the complexity and perfection of the cellular mechanisms, miniaturized to the utmost at the molecular level, which reveal within the cell an unparalleled knowledge of the laws of physics and chemistry
>
> — Albert Claude (1974)

> (iii) This attempt to isolate cell constituents might have been a failure if they had been destroyed by the relative brutality of the technique employed. But this did not happen.
>
> — Albert Claude (1974)

## 1. Introduction

The emergence of cellular life, both in the context of life’s historical origins and current research on ‘synthesizing life’, is one of the most complicated and least understood natural phenomena (Bernal, 1967; Lahav, 1999; Szostak et al., 2001; Luisi, 2006; Rassmusen et al., 2008; Gibson et al., 2010; Church & Regis, 2012). Arguably, the Big Bang about ∼13.6 billion years ago is better documented (by cosmic microwave background radiation) than the chemical and physical processes that can give rise to living cells.

The fact that living cells are non-equilibrium chemical systems suggests that life can emerge *only* from non-equilibrium chemical systems. Chemical non-equilibrium systems that continuously evolve arise naturally when solar irradiation strikes the surfaces of rotating planets (Spitzer & Poolman, 2009). The resulting cycling gradients of temperature, water activity, radiation, etc. drove planetary chemistries – the Earth’s ‘prebiotic soup’ of chemicals (Bada & Lazcano, 2003) – toward the contingent emergence of ‘primitive life’, and eventually toward progenotes and Last Universal Common Ancestors (Woese & Fox, 1977a, b; Woese 1998); at that point, about 3.5 billion years ago (Allwood et al., 2007), Earth’s chemical evolution became coexistent with biological evolution. The ‘bottom-up’ processes of prebiotic evolution can be investigated in chemical engineering simulators of Hadean Earth, with planetary sciences and geochemistry defining plausible initial conditions (Spitzer, 2013).

In this review, I take a complementary ‘top-down’ biological perspective and argue for experiments with cycling physicochemical gradients that act on a complex mixture of biomacromolecules obtained by killing a population of bacteria; the resulting ‘biotic soup’ contains all the ‘biochemistry’ needed to make living states emerge *de novo*. These experiments will clarify how the ‘architecture of living states’ – their spatiotemporal structuring – evolves from a mixture of ‘dead’ biomolecules and biomacromolecules (Harold, 2005).

As this approach reduces the complexity of life’s emergence to a well-defined though still very complicated *physicochemical* process, I will start – by way of an extended introduction – with general observations on the complexities of matter as differentiated by physics, chemistry and biology, on the irreducibility of biology to chemistry and physics, and on the transition from plausible speculations about life’s emergence to the new sciences of astrobiology and synthetic protocell biology (*cf*. Lord Kelvin counsel [i]). These introductory points expound the unique quandary of Darwinian (biological) evolution – the elusive nature of ‘life’s designers or drivers’: they are here rendered as persistently cycling physicochemical gradients.

### 1.1. The autonomy of physics, chemistry and biology

Theories and models of physics and chemistry have an extraordinary explanatory power derived from extensive use of high-level mathematics; in comparison, biology is generally perceived as having a different (less quantifiable) theoretical foundation (Mayr, 1996, 2007; Krakauer, 2011; Phillips *et al.*, 2009). Yet cytologists have known for some time that the living cell – the unit of biological matter – knows within itself the laws of physics and chemistry better than we can currently understand them; that there is nothing else within the cell but chemistry and physics – no teleological force to guide its evolution, no chance to play the tape of evolution again (Hinshelwood, 1946; Gould, 1989; *cf*. Albert Claude’s observation [ii]). We thus remain astonished at the physicochemical complexity of the cell, especially the spatiotemporal complexities of very small bacterial cells during their cell cycle – how they function as unique electrochemical systems converting environmental chemicals and energy into copies of themselves (Mitchell, 1979; Spitzer & Poolman, 2005, 2009, 2013; Spitzer, 2011). The question then is this: “How could simple (bacteria-like) cells come about *de novo* – now, in the laboratory, or at any time in the Universe?”

Physics explains inanimate matter in terms of partial differential equations, be they Newton’s equations of classical mechanics, Maxwell’s equations of electromagnetism, Schrödinger’s equation of quantum mechanics, or many other equations of mathematical physics. Such physical understandings, written down in the symbolic shorthand of advanced mathematics, were derived from simpler algebraic (phenomenological) laws, which summarized large amounts of numerical ‘regularities’ of experimental data (*cf*. Lord Kelvin’s comment [i], Krakauer, 2011). In physics, there are many such laws, such as Coulomb’s law of static electricity, Newton’s law of gravitation, Ohm’s and Kirchoff’s laws of electrical circuits, etc., which have been generalized into powerful theories of (mathematical) physics, often prized for their ‘beauty’ and internal self-consistency.

Chemistry has fewer differential equations, perhaps the most important being the equations of chemical thermodynamics and chemical kinetics; compared to physics, chemistry relies more on empirical laws and models (*e.g*. Faraday’s laws of electrochemistry, Guldberg-Waage’s law of mass action, Langmuir’s adsorption isotherm). Such chemical laws are sometimes understood through *ideal* (Platonic) models of *actual* matter, for example ideal solution, ideal conductor, ideal crystal and so forth, which more realistic models may approach under particular conditions. For example, the finite size of molecules and non-covalent attractive forces between them are accounted for in the van der Waals equation of real gases or the Debye-Hückel theory of electrolytes.

Biology has still fewer differential equations than chemistry; perhaps the best known is the Lotka-Volterra system of non-linear equations for prey-predator population dynamics. There are also fewer empirical laws, for example the statistical laws of Mendelian genetics, the Monod and other models of bacterial growth, and Michaelis-Menten enzyme kinetics (which is essentially a chemical model). The lesser degree of quantification in biology arises from the fact that living cells and organisms are too complex (or complicated) in relation to inanimate matter. Experiments with populations of bacterial cells are harder to quantify than those with ‘dead’ chemicals, because individual cells do not stay *chemically identical* as they increase in volume and split into two ‘imperfect’ copies – or in biologists’ language, ‘as they reproduce and evolve’; at cellular and subcellular levels, populations of bacterial cells are inherently heterogeneous (Neidhardt *et al.*, 1990; Kell *et al.*, 1991; Losick & Desplan, 2008). In comparison, a glycerol molecule for example, no matter by which chemical reaction it is (re)produced, is a rather constant and stable entity, carbon isotope effects notwithstanding.

Some aspects of cellular complexity can be understood when we consider the complexity of matter in general, going from physics to chemistry and from chemistry to biology. Physics deals with mass - ‘*everything’* has some mass (measured in kilograms); laws governing physical mass do not depend on its chemical composition, *e.g*. Newton’s laws of gravitation or viscosity. Chemistry differentiates mass as chemical mass with an *infinity of molecules and ions*, including charged and uncharged polymers, all made up from about 90 kinds of elemental atoms, and all having distinct atomic and molecular masses (measured in grams per mole). Biology differentiates chemical mass as ‘living’ mass made up from an *infinity of biological cells*, all of them (or most, at any rate) having genes encoded by nucleic acids (measured in base-pairs). The totality of genes defines the identity of the cell and guides its reproduction during the cell cycle in a given environment. Thus chemistry and biology are autonomous sciences (Mayr, 1996, 2007) dealing with their own ‘infinite kinds’ of complex matter, undergirded by the general laws of (mathematical) physics; however, where these sciences overlap, they provide a ‘deeper’, more fundamental (mathematical) understanding of the complexity of matter in general, no matter whether the initial impetus comes from physics, chemistry or biology.

Physics, particularly nuclear physics and quantum physics, clearly, has been astonishingly successful in explaining the origin, structure and properties of atoms (and hence the fundamental understanding of the periodic table and chemical bonds), and in theorizing about natural phenomena (*e.g*. interactions of electromagnetic radiation with matter that led to a multitude of spectroscopic methods, or the band theory of electrical conductors and semi-conductors that gave vacuum tubes, transistors, integrated circuits and sophisticated electronic devices). Physical instrumental techniques have proven immensely useful to chemists and biologists; X-ray diffraction in particular has become a key tool to determine the molecular structures of a multitude of polymeric materials, *e.g*. alumino-silicates (Pauling,), proteins (Perutz, Kendrew), and nucleic acids – the famous X-ray spots of Franklin’s DNA preparations that yielded the *chemical* explanation of biological heredity by Watson, Crick and Wilkins. However, physics cannot predict the *infinite* number of possible chemical molecules, particularly polymers whose molecular mass may become very large (approach ‘*infinity’*); physics cannot predict the stability of chemical compounds when exposed to different environmental conditions (*e.g*. water, temperature, irradiation), and physics cannot predict the methods of their syntheses (Krakauer, 2011).

Biochemistry, *i.e*. chemistry applied to compounds synthesized by biological cells, has determined the atomic compositions of biomolecules and biomacromolecules, their 3-D spatial structures including their chirality, their chemical and catalytic reactivities *in vitro* and their membranous, metabolic, and genetic functions. In particular, biochemistry has established: that all biochemical compounds have the same chemical properties, whether produced enzymatically by cells or by independent synthetic routes of organic chemistry; that all cells have the same but variable polymeric chemistry of nucleic acids, proteins, and carbohydrates, with just a few low molecular weight ‘building blocks’ and intermediates, including membrane-forming lipids and the ‘high-energy’ intermediates such as ATP and GTP; and biochemistry established that there exists a universal genetic code for all living cells and organisms and similar enzymatic mechanisms for regulating, transcribing and translating this code into proteins. Thus, there is a chemical unity to all life, and on that basis, molecular phylogenetics (Zukerkandl & Pauling, 1965; Woese, 1998; Pace *et al.*, 2012) suggests the historical existence of Last Universal Common Ancestors (LUCAs), when life began its Darwinian evolution on Earth about 3.5 billion years ago. However, biochemistry cannot predict the infinite number of possible biological cells and multicellular organisms, their appearances and dying-out as species, their interactions with variable environments and with each other, and the results of their continuing (non-equilibrium) evolution in general.

Taken together, the above considerations suggest that research at the ‘triple’ overlap of bacteriology, biochemistry and biophysics, could identify, at least operationally, the physicochemical mechanisms of how ‘living states’ come about from lifeless mixtures of biomolecules and biomacromolecules (Harold, 2005).

### 1.2. Speculation and facts: *Bathybius haeckelii*

Neither physics nor chemistry have shed much light on the physicochemical mechanisms of the cell cycle that a bacterial cell ‘knows’ so well, extending Albert Claude’s observation (ii) to bacterial cells, and which are embodied in Wirchov’s dictum *omnis cellula e cellula*. These ‘unknown’ physicochemical mechanisms are closely related to the ‘old’ questions of the emergence of cells *de novo* – the origin of life and its Darwinian evolution. Darwin was perhaps the first to suggest the emergence of life from lifeless *chemical* matter: he informally speculated that ‘primitive life’ might have emerged on early Earth in a ‘warm pond’ full of ammonium phosphates. (Remarkably, Darwin specified thermophilic conditions and singled out phosphate – the ionic form of phosphorus, which turned out as life’s crucial chemical element – for both metabolism and genetics).

How the issue of ‘spontaneous generation of life’ was put to rest by Pasteur’s dictum ‘*omne vivum ex vivo*’ is worth reiterating: Pasteur and his predecessors (*e.g*. Redi, Spallanzani and their ‘mistaken’ opponents) established qualitative (observational) *facts*, a pre-requisite to conduct quantitative and repeatable experiments – and then theorize about them (*cf*. Lord Kelvin’s counsel [i]). Thus Darwin, Pasteur and Wirchow regarded the question of life’s emergence *de novo* as too intractable to deal with, being gratified with the *factual* generalization: ‘all life only from life’. Thomas Huxley’s honest ‘mistake’ of regarding soft, gel-like inanimate matter as a new species of ‘primitive life’, *Bathybius haeckelii*, is noteworthy in this regard (Rice, 1983; Welch, 1984).

Even though our knowledge of life’s emergence *de novo* has remained speculative (Oparin, 1924; Bernal, 1967; Shapiro, 1986), such speculations do represent the beginning of scientific knowledge, which ought to lead to sound hypotheses and quantitative experimentation (*cf*. Lord Kelvin’s counsel [i]). Starting with Stanley Miller’s experiments (Miller, 1953) that demonstrated the synthesis of small biomolecules under pre-biological conditions (Bada & Lazcano, 2003), speculations about life’s emergence have now begun their transition into new biological sciences:

i. **Astrobiology**, which deals with the *historical* issues of the emergence of first cells (Gilmore & Sephton, 2004; Sullivan *et al.*, 2007; Lahav, 1999; Fry, 2000; Deamer, 2011); the primary drivers for their emergence were persistently cycling, chemical disequilibria arising from solar irradiations of rotating planetary surfaces (Rothschild, 2003; Spitzer & Poolman, 2009; Spitzer, 2013; Stüeken *et al.*, 2013); and
ii. **(Semi)synthetic protocell biology**, which deals with *contemporary* laboratory constructions of protocells (vesicles) that enclose designed systems of genetic circuits and their metabolic and regulatory components (Szostak *et al.*, 2001; Luisi, 2006; Solé, 2007; Mann, 2012; Rassmusen *et al.*, 2008; Nandagopal & Elowitz, 2011; Purnick & Weiss, 2009; Kwok, 2010; Noireaux *et al.*, 2011; Schwille, 2011; Benner & Sismour, 2005; Church & Regis, 2012; Gibson *et al.*, 2010; Elowitz & Lim, 2010). Neither of these approaches has yet demonstrated a plausible emergence or construction of cellular life *de novo*, the inherent technical challenges being compounded by the question ‘What is life?’ (Pross, 2011, 2013; Benner, 2010; Trifonov, 2012; Szostak; 2012; Spitzer, 2013).

### 1.3. Conceptual quandary: the ‘designers’

From a physicochemical standpoint, a complex bacterial cell cannot self-(re)assemble *spontaneously* (via attractive non-covalent molecular forces) without the action of external agents or ‘designers’ – be they natural (unintentional), *e.g*. Earth’s dynamic diurnal environments (Spitzer & Poolman, 2009; Spitzer, 2013), or anthropic (intentional) such as the brain’s capacity to ‘design and build’ (Szostak *et al.*, 2001; Gibson *et al.*, 2010; Elowitz & Lim, 2010) – or yet to be discovered.

This conceptual quandary suggests an experimental approach where the driving force for biological self-assembly and evolution are cycling gradients of temperature, water activity and electromagnetic irradiation acting on mixtures of dead biomolecules and biomacromolecules; specifically, I suggest ‘making life emerge *de novo*’ from a lifeless mixture of *all* biochemicals of which a bacterial cell is composed. Such a biochemical mixture – a ‘dead (unstructured) biotic soup’ (Harold, 2005) – can be obtained by *gently* killing a living bacterial population (*cf*. Albert Claude’s observation [iii] about the *brutality* of killing eukaryotic cells). I briefly discuss how to make biotic soups and recover ‘living states’ from them by applying physicochemical gradients. I conclude that there exists *cycling non-equilibrium continuity* between ‘life’ and ‘non-life’, *i.e*. being alive and reproducing, or being dead (Davey, 2011).

## 2. Why do bacterial cells appear so complex?

I take a simplifying physicochemical approach to understanding (biological, bacterial) complexity, which has been described in terms such as: emergent behavior, self-assembly, non-linearity, pattern-formation, dissipative self-organized structures, operating ‘on the edge of chaos’, fragile yet robust, hierarchical networks, feedback loops, system control, catastrophic cascades, autocatalytic cycles of ‘replicators’, ‘order out of chaos’, biosystems, etc. (Adami *et al.* 2000; Goldenfeld & Kadanoff, 1999; Schliwa, 2002; Mazzocchi, 2008; Krakauer, 2011; Csete & Doyle, 2002; Hazen, 2005; Van Regenmortel, 2004; Bertalanffy, 1950; Qian, 2011; Kell & Welch, 1991; Whitesides & Ismagilov, 1999). Some of these concepts, borrowed from network and process control engineering, are realized in solid-state electronic devices of telecommunication and chemical manufacturing processes, including computer-controlled sensors and regulators.

Disregarding the issue of ‘design and designers’ for a moment, the chemical basis of electronic devices lies in the semi-conducting properties of doped silicon and similar materials. The obvious difference between the solid-state operation of electronic ‘chips’ and the functioning of biological cells is the soft, semi-liquid (partly gelled) nature of cells (Bray, 2005, 2009; Fels *et al.*, 2009; Spitzer & Poolman, 2013), which allows for the possibility of their *natural self-reproduction*: two bacterial cells arise from one by about 1500 kinds of parallel and sequential biochemical reactions, sometimes in less than 30 minutes if external environment (nutrient medium) is favorable; ‘life’ in the solid state only (no molecular translational motion) where molecules ‘sit’ is harder to imagine (http://schaechter.asmblog.org/schaechter/2013/03/feynman-said-just-look-at-the-thing-.html). Biological cells are therefore materially and functionally far more complicated than electronic computers to which they are sometimes likened (Danchin, 2009). Such electronic devices operate in the absence of chemical reactions in *spatiotemporally fixed designs* of semi-conducting silicon chips. Still, as a common denominator, both (designed) solid-state electronic devices and (non-designed, evolving) ‘soft’ biological cells conduct *electrical charges* - but of vastly different kinds and by vastly different mechanisms - and generate self-regulated electrical potentials. In that sense, bacterial cells have a ‘brain function’ that has the same physical basis as computers and other biological cells (particularly cells of nerve tissues): the storage and movement of electrical charges, including their ‘creation’ and disappearance by chemical reactions.

The physicochemical complexity of bacterial cells lies in several interdependent characteristics: (i) ionic and multicomponent, (ii) multiphase, (iii) ‘crowded’ (highly concentrated), and (iv) self-regulated and re-emergent. The first three characteristics define ‘complex chemical matter’; it can be either *equilibrated* (‘definitely dead’), or *kinetically arrested* away from equilibrium (‘dead’ but able to resume its chemical reactions in the future); or it can be in a *steady state*, chemically reacting but not evolving, its properties being independent of time; or it can be in a *non-steady state*, chemically reacting and *evolving* in time – the pre-requisite state of complex chemical matter from which living states can emerge and evolve. The fourth characteristic is quintessentially biological, as *self-regulation of re-emergence* requires the cyclic maintenance of chemical and physical identity – in biological language, the maintenance and continuation of the genome (nucleoid, plasmids, and even phages) and of the membrane (cell envelope) during the cell cycle. These four elements are further discussed below.

### 2.1. Ionic and Multicomponent

Bacterial cells are composed of many kinds of chemical compounds, of a very wide range of ionic character, molecular weights, and hydrophobicity, from small ions and molecules in the sub-nanometer range to ‘enormous’ polyelectrolytes (nucleic acids) in the micrometer range; a great deal of biochemistry has been concerned with their isolation and purification, and with determinations of their chemical compositions, 3-D chemical structures, *in vitro* biochemical reactivity and self-assembly, and their cellular localizations and physiological functions. There are relatively few strictly *uncharged* biomolecules (and no biomacromolecules) in a bacterial cell, except for ‘special cases’ of amphoteric compounds like proteins at their isoelectric points, or zwitter-ionic lipids and similar compounds derived from choline. Uncharged ‘food’ molecules are normally available in ionic form, *e.g*. carbon dioxide as hydrogen carbonate ion, or they are first converted by kinases to anionic organic phosphates in order to become available to cell’s biochemistry.

In a bacterial cell, there are over 1000–2000 kinds of proteins, many kinds of RNAs, plus a single DNA polymer, and 3-6 kinds of phospholipids; there are also ‘inorganic’ chemicals of non-biological origin that are important reactants or catalysts (widely available from the environment), chief being water, carbon dioxide, nitrogen and oxygen, and inorganic ions, importantly potassium, sodium, magnesium, calcium, iron, bicarbonate, mono- and di-hydrogen phosphates and sulfate and chloride; multivalent cations (Fe, Mg, Ca) are normally sequestered in ‘organic’ molecules and biomolecules, their free concentrations being extremely low. Physicochemical interactions between bacterial biomolecules and biomacromolecules enable the functioning of cellular biological mass, mediated by water and simple inorganic ions. The effects of ions are sometimes manifested in an unexpected manner, *e.g*. the initiation of embryological development of unfertilized sea urchin eggs by a specific solution of mixed salts (Loeb, 1899, 1906; Pauly, 1987). Water molecules and simple ions participate in many biochemical reaction pathways (synthetic, degradative and signaling), and in this *chemical* sense cellular life depends critically on *aqueous* (multicomponent) electrolyte (Spitzer & Poolman, 2005; Spitzer, 2011). There is likely no biochemical reaction that does not feature water or some ‘aqueous ionic species’, either as reactants or products, or catalysts.

These ‘rough’ compositions, discussed in detail in current textbooks (Neidhardt, *et al.*, 1990, White, 2000; Phillips *et al.*, 2008; Schaeter *et al.*, 2006; Kim & Gadd, 2008), support the contention that a bacterial cell operates within an electrochemical paradigm that includes ‘battery-like’ electrolytic conduction (Mitchell, 1979), because essentially all macromolecular components (the nucleoid, protein-protein complexes, protein-RNA complexes and their clusters) are charged and dissolved or dispersed in cytoplasmic water of relatively high ionic strength; changes of ionic strength of external environments are converted to localized ionic changes within the cytoplasm, activating constitutive membrane proteins or genetic circuits within the nucleoid (Higgins *et al.*, 1987; van der Heide *et al.*, 2001). The bacterial membrane is also ionic – it contains anionic lipids (Yeung *et al.*, 2008), including highly charged cardiolipin (Koppelman *et al.*, 2001; Romantsov *et al.*, 2008) and ‘semi-conducting’ membrane proteins. Finally, as viewed from the nutrient medium, the bacterial cell *in toto* is electrically charged, in a complex and patchy manner, as evidenced by cells’ electrophoretic motion in an external electric field (van Loosdrecht *et al.*, 1987; Bayer & Sloyer, 1990; de Kerchove & Elimelech, 2005).

### 2.2. Multiphase

On account of the physicochemical variety of bacterial biomacromolecules (charged, hydrophobic, hydrophilic), *attractive* non-covalent molecular forces bring about multicomponent ‘phase separations’. The attractive forces responsible for the existence of liquid phases in general are van der Waals forces and ion-dipole and similar ‘Debye’ type polar forces; however, the most important ‘cytological’ non-covalent forces are hydrogen bonding (Pauling *et al.* 1951; Eisenberg, 2003), the hydrophobic effect (Tanford, 1980; Southall *et al.*, 2002), and (screened) electrostatic attractions between cationic and anionic biomacromolecules (Schreiber & Fersht, 1996; Halford, 2009). The crucial role of electrostatic attractions is also manifested by the bimodal distribution of isoelectric points of proteins (Spitzer & Poolman, 2009), with peaks in the alkaline pH region (negatively charged proteins) and acidic region (positively charged proteins); such bimodal (‘positive-negative’) distributions are common across many proteomes (Kiraga *et al.*, 2007), suggesting that electrostatically assisted protein clustering (gelation) is a general cytoplasmic phenomenon of many kinds of cells. The effects of attractive forces are modulated by repulsive non-covalent forces, in particular by the excluded volume effect (Zimmermann & Minton, 1993), *i.e*. by the hydration stabilization of ‘folded’ macromolecules in water (Halle, 2004; Persson & Halle, 2008; Jasnin, 2009; Fenn *et al.*, 2009) and by screened electrostatic repulsions between anionic molecular surfaces (Spitzer, 1984; 2003; Poolman *et al.*, 2004; Spitzer & Poolman, 2005).

The complexity of this physicochemical situation precludes any chances that a cell could ‘self-assemble’ *spontaneously* from its components, as shown by a simple Gedankenexperiment: in a large beaker of water, let us separate all the biomacromolecules and biomolecules and ions of a cell to infinity from each other and let then water evaporate from this ‘infinitely dilute bacterial broth’. As the water evaporates, attractive non-covalent forces become operative and bring about phase separations; however, it is quite unlikely that a cell would re-form and ‘start living’ just by letting all the molecular surfaces come into close proximity when water evaporates. We are quite sure that randomly agglomerated and gelled materials will sequentially phase-separate as ‘dead’ chemicals (Walter & Brooks, 1995; Stradner *et al.*, 2004; Kayitmazer *et al.*, 2007). Thus the existence and operation of (fundamental) *non-covalent* intermolecular forces cannot explain how a cell could self-assemble from its components, even though such forces are necessary (but not sufficient) to drive the formation of the membrane and of other biomacromolecular structures.

Other factors are at play, the clue being provided by bacterial physiology: the dynamic architecture of a bacterial cell *in vivo* depends on the *rate* of its growth (Neidhardt *et al.*, 1990; Scott *et al.*, 2010; Wagner *et al.*, 2009), *i.e*. on the rate of synthesis and degradation of its components, like DNA, proteins, RNAs, ribosomes and its membrane (lipids and membrane proteins). It is on this dynamic process of biochemical reactions that the molecular non-covalent forces act, bringing about particular, time-dependent and localized phase separations of supramacromolecular structures – the dynamic architecture of the cell (Harold, 2005); this dynamic architecture is lost when cells are taken apart for analyses of their biochemical components.

Phase separations of any kind imply classical physicochemical concepts of high concentrations of partially miscible (or immiscible) systems and saturated solutions, and it has long been argued, both from biochemical and physicochemical standpoints (Ovadi & Srere, 1999; Zhou *et al.*, 2008) that *in vivo ‘*crowding’ is an important fact, the consequences of which have been somewhat neglected (Ellis, 2001).

### 2.3. ‘Crowded and Supercrowded’

On a micrometer scale and on the seconds or minutes timescales, a textbook picture of a bacterial cell can be regarded as a system of four physicochemical ‘phases’ (each being also multicomponent): (i) the hydrophobic cell envelope of a particular geometrical shape (coccoid, rod etc.), the key functional part being a lipid bilayer studded with a high number of many kinds of ‘water-insoluble’ proteins, (ii) the phase-separated nucleoid, portrayed like a bundle of ‘spaghetti’ representing anionic DNA helices condensed by cationic proteins and amines, (iii) the ribosomes, phase-separated clusters of proteins and nucleic acids (‘water *and* lipid insoluble’), shown as tiny spheres scattered throughout the cytosol (Alberts *et al.*, 2002), or attached to mRNA strands (polyribosomes) that are connected to DNA through RNA polymerase (Schaechter *et al.*, 2006), and (iv) ‘unstructured’ cytoplasm, an aqueous solution or dispersion of proteins, nucleic acids, metabolites and ions. On this micrometer scale, cytoplasmic proteins are not shown because they are too small; at greater spatiotemporal resolutions, proteins and their (hetero)dimers and multimers (complexes) are shown ‘randomly crowded’ (Goodsell, 2010), representing ∼ 25% of the volume of the cytoplasm (Zimmerman & Trach, 1991). An important intracellular phenomenon is polymer incompatibility of hydrated nucleoids, plasmids and phages (mostly nucleic acids) with crowded cytoplasmic proteins (Valkenburg & Woldringh, 1984; Woldringh & Nanninga, 2006); this represents a major large-scale phase separation (compartmentation, or biomacromolecular localization) within the bacterial cytoplasm, a precursor phenomenon to the formation of the fully compartmentalized nucleus of eukaryotic cells.

Zooming down to nanometer scale and microseconds and longer timescales, biomacromolecular surfaces can be considered as smoothed-out, or averaged out, into four kinds of distinct physicochemical patches (Wang *et al.*, 2011; Spitzer, 2011): hydrophilic (capable of hydrogen bonding with water molecules), hydrophobic (unable to be accepted into the hydrogen-bonded network of water molecules), and positively or negatively charged (Poolman *et al.*, 2004). These ‘smooth’ chemical patches give rise to repulsive and attractive forces that determine to what degree a biomacromolecule is attracted to and oriented vis-à-vis its neighbors, *i.e*. its clustering propensity with other biomacromolecules, which can result in the formation of a gel phase of variable mechanical (viscoelastic) strength.

Simple physical models of such high crowding (Phillips *et al.* 2009; Spitzer & Poolman, 2009, Goodsell, 2010; Spitzer 2011) show that cytoplasmic biomacromolecules are essentially ‘touching’ *in vivo*, and thereby interacting strongly via repulsive non-covalent molecular forces (hydration and screened electrostatic repulsions) and thus avoiding catastrophic (fatal) large-scale agglomerations or precipitations (Pastore and Temussi, 2011). The functional clustering of biomacromolecules has been also theorized by the concept of metabolons, where reactants and products are shuttled between the catalytic sites within the cluster without bulk diffusion in the cytoplasm (Srere, 1985; Mathews, 1993, Ovadi & Srere, 1999), as modular hyperstructures (Norris *et al.*, 2007; Hartwell *et al.* 1999), and as various ‘omes of system biology, such as FtsZ divisome (Lutkenhaus *et al.* 2012) which is synthesized and assembled before cell division takes place; some ‘omes’ have now been rendered visible (Bakshi *et al.*, 2012; Fu *et al.*, 2010; Greenfield *et al.*, 2009).

Within the supramacromolecular ‘supercrowded clusters’, also described as a ‘sieve-like meshwork’ of cytoskeleton and other proteins (Mika *et al.* 2010) or a ‘poroelastic network’ in the case of eukaryotic cells (Charras *et al.*, 2008), the biomacromolecules are separated by narrow channels filled with percolating multi-ionic electrolyte solution that hydrates (stabilizes) biomacromolecular surfaces and thus maintains their biochemical and ‘architectural’ functions (Harold, 2005). Such channels are theorized to have semi-conducting electrolytic properties, *i.e*. the capability to sort out and direct both inorganic ions and biochemical ions into different parts of the cell. These channels could be considered as transient ‘microfluidic *electrolytic* integrated circuits’,ultimately ‘programmed’ by the nucleoid (genome). The semi-conductivity of these channels is based on the solution of a system of differential Poisson-Boltzmann equations applied to interacting planar anionic surfaces with an electrolyte in between them; under certain conditions – high negative potentials at a given temperature for a given set of charges, the electrolyte space between the surfaces becomes conductive only to cations, with anions being prevented to enter the space between the anionic surfaces (Spitzer, 1984; Spitzer, 2003; Spitzer and Poolman, 2005). Phosphorylation reactions, which increase the negative potential within the channels, thus act as ‘switches’ that direct and sort out anions (according to their charge), *e.g*. ATP, ADP etc., toward more positive potential regions within the supercrowded clusters; at the same time, they change the morphology of the clusters by increased screened electrostatic repulsions (Spitzer, 1984; 2003).

The crowding *in vivo* can significantly increase, up to ∼50% volume fraction, in hyperosmotic media, when the cell loses water; protein sensors and transporters within the cell envelope then become activated and import osmolytes (potassium ions, glycine betaine, trehalose, etc.) in order to reduce the osmotic outflow of water into the nutrient medium (van den Bogaart *et al.*, 2007; Morbach & Krämer, 2002; Wood, 2011), and thus render plasmolysis less severe. Astonishingly, *Escherichia coli* cells can re-adjust and resume growing, albeit more slowly, under such extreme ‘supercrowded’ (hyperosmotic) conditions (Konopka *et al.* 2009), provided that the increase of crowding is performed in a step-wise sequential manner, and the cells have time to restructure its spatiotemporal architecture – to adapt to higher crowding conditions, when there is just about a monolayer of water of hydration around biomacromolecular surfaces (Cayley *et al.*, 1991); however not all cells appear to survive in such experiments, reflecting the heterogeneous nature of bacterial (biological) populations.

The large concentration of cytoplasmic hydrated biomacromolecules has been theoretically treated as the repulsive excluded volume effect, which predicts increased stabilization of protein structures (Zhuo *et al.*, 2008) and increased tendency to form biomacromolecular complexes – various (hetero)multimers of proteins or proteins with nucleic acids; this effect has been investigated *in vitro* with polymeric ‘crowders’ (*e.g*. dextrans, polyethyleneglycols, etc.) as model polymers to simulate the crowding inside living cells. While this approach has been partly successful, it has now become apparent that *attractive* non-covalent forces between biomacromolecules need to be also considered: there is a ‘thin line’ between biochemically regulated clustering (gelling) and non-functional (fatal, irreversible) aggregation of proteins and nucleic acids (Pastore and Temussi, 2011; Jiao, *et al.*, 2010; Ellis & Minton, 2006; Miklos *et al.* 2010; Wang *et al.*, 2011; Harding & Rowe, 2010; Schlesinger *et al.*, 2011; Boquist & Gröbner, 2007; Gierasch & Gersheson, 2009).

### 2.4. Self-regulating and re-emergent

The above description of the physicochemical complexity of a bacterial cell – ionic and multicomponent, multiphase, and crowded, is made more complicated by the fact that during the cell cycle there are about 1000–2000 concurrent and sequential biochemical reactions within the cell and between the cell and the environment; these biochemical reactions are rather well-synchronized and regulated to yield physiological processes, such as: (i) the sensing of the extracellular environment and the import of nutrients into the cell by the cell envelope, (ii) the conversion of the extracellular signals into cytoplasmic signals and their reception by the cytoplasmic side of the membrane and by the nucleoid (iii) biosynthesis of low molecular weight ‘building blocks’ including ‘fueling molecules’ such as ATP and GTP (iv) activation/deactivation of constitutive membrane proteins for immediate responses to environmental inputs, (iv) gene activation and silencing, and transcription, (iv) biosynthesis of ribosomes (v) translation via ribosomes, (vi) the initiation, control and termination of the enzymatic replication of the nucleoid and plasmids, and (vii) the cell division and other morphological movements (shrinkage, invagination, budding, adhesive gliding, sporulation, etc.)

There can be little doubt that coulombic (electrostatic) interactions and electrolytic semi-conduction play a major role in regulating such complex processes on a global cellular scale; this is so because ‘naked’ electrostatic forces (Coulomb’s law, eq. 1) are both very *strong and long-range* compared to *all* other non-covalent molecular forces.

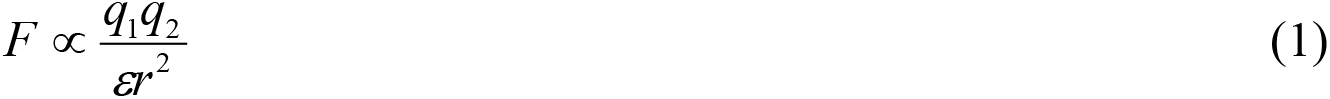

Here, the force *F* is inversely proportional to the dielectric constant *ε* of the solvent and to the square of the distance *r* separating the charges *q1* and *q2*. The cell *must* (and does) operate in an aqueous electrolyte of a relatively high ionic strength in order to shorten the range of naked coulombic forces (eq. 1), and thus make them *commensurate* with other non-covalent molecular forces (especially hydration) on the scale of a little below one nanometer (Spitzer & Poolman, 2009). The Debye-Hückel theory then modifies Coulomb’s law, eq. (1) approximately by the exponential term as

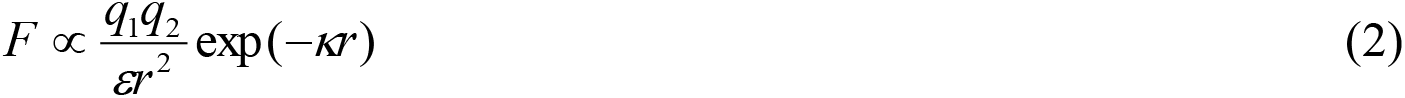

Ionic strength is proportional to the square of the Debye constant *κ*, the inverse of which *1/κ*, is a measure of the effective range of screened electrostatic interactions: the higher the ionic strength, the faster the screened electrostatic forces decay. The high dielectric constant *ε* of water *reduces* the magnitude of electrostatic forces over any range of distances, be they ‘naked’ or screened (Phillips *et al.*, 2009; Spitzer & Poolman, 2009); the high dielectric constant of water and its unique temperature dependence thus selects water as the biological solvent – there are no other solvents with similar or higher dielectric constants so readily available, providing an environment ‘fit for life’ (Henderson, 1913; Ball, 2008; Poolman & Spitzer, 2009). The commensuration of screened electrostatic forces, hydration and high biomacromolecular crowding suggests that the well-established 2-D membrane vectorial biochemistry is extended into 3-D vectorial biochemistry within the cytoplasm, particularly on the cytoplasmic side of the membrane (Spitzer, 2003; Poolman et al., 2004; Spitzer & Poolman, 2005, 2009; Spitzer 2011; Spitzer & Poolman, 2013).

It is not coincidental that when a bacterial cell is stressed (*e.g*. by starvation), the electrostatically highly charged molecules of (p)ppGpp are synthesized in order to ‘alarm the cell ‘electrically’, ringing the doorbell so to speak, which initiates the *stringent survival response* on a global cellular scale; such response includes the re-switching of genetic circuits, progressive shutting-down of transcription and ribosome biosynthesis, the densification of the nucleoid, the shrinking of the cell volume, etc. (Neidhardt *et al.*, 1990; Potrykus and Cashel, 2008; Siegele & Kolter, 1992; Frenkiel-Krispin *et al.*, 2004).

Fundamentally, a bacterial cell, at any instant of its growth, can be viewed as a system of biochemically reacting carriers of electrical charge: charged proteins and nucleic acids dissolved and dispersed in an aqueous electrolytic solution, separated from the environment by a charged lipid bi-layer containing anionic lipids and ‘semi-conducting’ protein transporters and channels (and a host of other proteins); localized biochemical reactions then ‘charge and discharge’ the membrane, which acts as a (‘leaky’) electrical *viscoelastic capacitor*, maintaining variable membrane potential and creating a multitude of chemiosmotic signals on the cytoplasmic side of the cell envelope (Mitchell, 1979). Some of these signals are then transmitted ‘tangentially’ along the cell envelope to control for example the motion of the flagellum, and some are transmitted to the nucleoid to regulate gene expression, and thereby cell’s growth, manifested by bio-electromechanical (morphological) movements of the cell (Morris & Jensen, 2008).

In order for such a complex system to work reasonably reproducibly during the cell cycle (as it does), biochemical reactions have to be localized within the cell and within the cell envelope. A relatively simple physicochemical mechanism to localize biomacromolecules is for them to become incorporated in 2-D ‘rafts’ within the cell envelope (Bray, 2005; López & Kolter, 2010) and in ‘supercrowded’ 3-D gels within the cytoplasm, as hypothesized by the sol-gel model (Spitzer & Poolman, 2013). In this model, ‘supercrowded’ biomacromolecular clusters (gels) are associated mainly with the cytoplasmic side of the cell envelope, and the nucleoid is situated in the middle, though many other spatiotemporal arrangements are possible; a similar interpretation was given to protein diffusion data in mitochondria (Partikian *et al.*, 1998).

The biomacromolecular gel formation (supercrowding), and the reverse process of gel liquefaction, are controlled by biochemical signaling reactions that increase or decrease hydrophobic and screened electrostatic attractive interactions (*i.e*. epigenetic modifications or ‘processing’ of nucleic acids and proteins during or after their biosynthesis); for example, methylations and dephosphorylations increase non-covalent attractions (gel formation) and the reverse reactions increase repulsions (gel liquefaction). Such regulated sol-gel transitions contribute to the morphological changes of the cell, along with the forces generated by the electrical charging and discharging of the lipid bilayer membrane (viscoelastic capacitor).

## 3. Experiments with ‘bacterial biotic soup’

As the above descriptions show, any bacterial cell cycle is a very complicated process taking place under molecularly crowded conditions with myriads of physicochemical interactions. The molecular details of these interactions are currently being untangled by the approaches of system and synthetic biology – the ‘furry balls’ of interactomes, transcriptomes and other ‘-omes’, and the ‘dumpling soups’ of signaling pathways (Lewitzky *et al.*, 2012; Tompa & Rose, 2011), as well as by computational approaches and modeling of the whole cell or its particular physiological functions (Elcock, 2010; Cossins *et al.*, 2011; Goldman et al., 2009).

To cut through such formidable complications when considering the living cell *in toto*, I suggest an experimental approach – a short-cut in effect, to reconstructing ‘life’, *i.e*. a general, open-ended experimentation with ‘biotic soups’, *cf*. Introduction. The biotic soup experimentation would take place in two stages; first, the making of ‘biotic soups’, and second, subjecting them to cyclic gradients of temperature, water activity, radiation (UV, visible) etc., echoing the cyclic temperature gradients of the polymerase chain reaction (PCR). The cycling gradients also simulate Earth’s diurnal cycling as the primal force that brought about the historical emergence of cellular life and has contingently driven its evolution since (Rothchild, 2003, 2008; Spitzer & Poolman, 2009; Spitzer, 2013), as evidenced by the circadian clocks of many types of cells, including bacterial cells (Wijnen & Young, 2006). I thus argue for: (i) development of laboratory protocols to grow and disassemble bacterial cells (by physical means) from a ‘living’ state to a ‘dead’ state, and (ii) development of protocols to recover ‘living states’ from such disassembled bacterial parts, principally by cycling gradients of temperature, water activity, with a judicious ‘feeding’ of nutrients.

This approach requires expertise in growing, characterizing and manipulating living and dead bacterial populations, hence I discuss this topic only briefly in the form of questions, which bacteriologists and biotechnologists can address with the requisite knowledge in order to develop more detailed experimental designs.

### 3.1. Making ‘biotic soup’

Any bacterial population, possibly in a biofilm, can be lysed, yielding a lysate or a dead ‘biotic soup’ for further characterization and experimentation; the ‘biotic soup provides all the necessary chemicals more or less in the right proportions from which a ‘living state’ could arise, not necessarily in the form or growth habit of the original bacterial species. The following questions need be carefully considered:

i. *Bacterial species*. Which bacterial populations (species, strains, metabolic life-styles, etc.) to select: ‘minimal’ gene and physical size, or a specific viable strain, L-forms (Leaver *et al.*, 2009; Diennes & Sharp, 1956), thermophiles, photon-dependent autotrophs? Non-oxygen respiring cyanobacteria may provide the most suitable model in relation to the origin of life questions, being able to grow without complex organic molecules, and being naturally transformable?
ii. *Growth conditions*. Which nutritional medium to select or design? Making this choice is closely related to the above question; some minimal medium would be the most suitable from the standpoint of the origin of life research, though not necessarily from the point of finding the physicochemical ‘laws’ that describe the conditions of life’s emergence.
iii. *The method of killing*. How to terminate a living bacterial population? How to lyse a bacterial population? A method needs be designed in which no ‘unnatural’ chemicals, such as surfactants, solvents or enzymes are added; physical means are preferred, such as the lack of specific nutrients (starvation), a mild hypo-osmotic shock, various degrees of sonication, and different rates of freezing in a variety of nutrient media of different water activities, slow evaporation of water, and any combination of such approaches.
iv. *Timing of killing*. At which point in the growth curve to ‘kill’? Mid-point exponential, early on in the population growth with sufficient ‘food’ left over, or a stationary stage, or some particular state of nutritional or environmental stress?
v. *Is the biotic soup dead?* What would be the assay for making sure that the biotic soup is ‘really dead’? (Davey, 2011). Not an easy experimental question, though current microbiological protocols would be a good starting point. [An aside: Would we conclude that a biotic soup was ‘primitively alive’, if a virus or phage were to multiply in it under cycling temperature conditions?]
vi. *What would be the supramacromolecular composition of the biotic soup?* Most likely, there will be ribosomes, plasmids and nucleoids, all partly broken, with partly attached ‘broken’ cell envelopes (Tremblay *et al.*, 1969; Firshein, 1989; Zimmerman & Murphy, 2001); their kinds and concentrations will depend on the particular selections above.

The composition of biotic soups is clearly very complex and new characterization methods will need be developed, for example ‘gentle’ centrifugal fractionation to obtain new kinds of ‘less complex’ fractions. Because extant bacterial populations are biochemically robust with redundant systems to survive, some simplification of biotic soups may be possible and in fact desirable in order to better understand the essential principles of ‘recoverable’ living states.

### 3.2. ‘Putting Humpty-Dumpty together again’

According to the notion that Earth’s diurnal cycles, yielding a multitude of cyclic physicochemical gradients, were in effect the ‘designers’ of living systems, such gradients may also perform ‘repeat’ experiments with (simplified) biotic soups in continuous (evolutionary) attempts to re-construct some form of a living system. Thus, in the temperature upswings the structures will tend to diminish (dissolve and/or dissociate more, though not necessarily), and in the temperature downswing, they might re-appear, perhaps in a different manifestation (structural details), given the supramacromolecular nature of the system. Also, the cycling temperature regime provides energy when the temperature goes up; this mechanism speeds up chemical reactions, and also plays a role in coupling signaling reactions and sol-gel phase transitions.

Thus cyclic temperature gradients both *shape* supramacromolecular colloidal structures needed for ‘life’ (through non-covalent attractive and repulsive interactions inherent in biomacromolecular crowding) and *drive* some biochemical reactions more than others; in other words, the stoked biotic soup may ‘live’ when the temperature goes up, and become dormant or dead when the temperature goes down. This model could represent Darwin’s primitive ‘semi-life’, dependent directly on external ‘heat stoking’ metabolism, though such a semi-life may not be readily recognizable by current biochemical assays.

As with the preparation of the biotic soup, there are a number of questions to consider regarding the ‘stoking’ regimes:

i. What would be the starting volume fraction of all biomolecules and biomacromolecules? We could start with a very dilute system as in *in vitro* biochemistry or with a very crowded (*in vivo* range of crowdedness) or even with a supercrowded system; this will depend on the lysing methods and other assay methods. Thus the ‘biotic soup’ could be stoked in an open evaporating system in order to bring it to the *emergent crowding transition* (Spitzer and Poolman, 2009), or it could be stoked supercrowded at, say, a 70% volume fraction while judiciously adding water and nutrients (Spitzer & Poolman, 2013); alternatively, if the final system is in the range of *in vivo* volume fractions (20–50%), then a closed system with constant volume fraction might be in order.
ii. How to design the reactor as an ‘evolving chemostat’? How to synchronize feeding regimes of nutrients with cooling/heating cycles? How to determine biochemical conversions (the ‘growth of the biotic soup’) and ‘define’ operationally a living state?
iii. How would the final ‘structure’ of such a biotic soup (in the 30–50% range of volume fractions) depend on the frequency and amplitude of the temperature cycles? Again, these choices will have to be made in the light of the biochemical suitability of bacterial species to withstand large temperature changes without denaturing the functional conformations of key biomacromolecules; hyperthermophiles may be a suitable choice here.
iv. How could we detect the formation of transient supramacromolecular structures, and membranes with integral proteins, and their biochemical activity?
v. How could we monitor the structural state of the chromosome, and its replicating activity?
vi. What effects could be observed by increasing the ATP/ADP ratio? It is hypothesized that increasing the ATP/ADP ratio could ‘jump-start’ transient reaction loops, when the biotic soup reaches the ‘emergent’ level of biomacromolecular crowding; when to start ‘feeding’ the emerging living state?
vii. What would be the criteria to distinguish between the living, comatose, sick or injured, dormant, viable but non-culturable, and dead?

Any emergent living state may not readily materialize in the classical prokaryotic physical forms (coccoid, rod etc.) because the formation of a *closed cell envelope with a lipid bilayer membrane* is expected to be the most challenging phenomenon to bring about and manipulate by cycling physicochemical gradients: most likely, the cycling experiments need to be done in the ‘supercrowded’ regime (Spitzer, 2011; Spitzer & Poolman, 2013), when ‘super-cells’ (with imperfect internal compartments and multiple chromosomes, in effect ‘proto-biofilms’) will initially start assembling and disassembling. Such unstable pre-Darwinian ‘barely living states’ may be identified as progenotic, *i.e*. without established heredity and organismal lineages – with rampant horizontal gene transfer – but with a stable (no longer evolving) genetic code (Woese & Fox, 1977; Fox 2011; Fox *et al.*, 2012).

## 4. Bacterial cells as evolving chemical micro-reactors

The model of a sol-gel, battery-like workings of a cell as a ‘*viscoelastic capacitor*’ is consistent with a more general thermodynamic view of a growing cell as an open thermodynamic system that exchanges energy and materials with the surroundings in accordance with the laws of chemistry and physics (Hinshelwood, 1946; von Bertalanffy, 1950; Qian, 2006). In practice, such open non-equilibrium thermodynamic systems are industrial chemical reactors, and it is therefore useful to compare the operation and regulation of a chemical reactor with that of a bacterial cell. This comparison has its limitations, just as comparing a cell to the hardware and software of a computer can be stretched only so far; still, such comparisons do illuminate the workings of a cell as a complex self-programmed chemical engineering process, which, as any such process, is prone to stochastic and contingent errors, malfunction and even catastrophic upsets.

### 4.1. Ideal bacterial cell

Admittedly, a bacterial cell is a very peculiar chemical micro-reactor; compared to an industrial chemical reactor; there are two obvious differences: (i) the walls of a chemical reactor have fixed shape and stay constant and are not part of the chemical production as compared to the cell envelope (however they are ‘permeable’ as raw materials go in and the product gets out in a regulated and prescribed manner), and (ii) the solid-state computer with an appropriate program that regulates and controls the reactor resides outside and is not part of the production either, as compared to the ‘nucleoid program’ that is inside the cell. In both cases there are many sensors and regulators that monitor and control the process inside and outside.

The operation and regulation of this unusual chemical micro-reactor can be elaborated with the help of an ‘ideal biological cell’, similar to the concept of ideal solution of physical chemistry. The ideal biological cell has two characteristics: a) its operation and regulation (growth and division) are without errors, and b) its environment is ‘infinite’ and unchanging. Under such ideal conditions, only ‘mathematically’ exact, non-evolving copies of the ‘mother cell’ will be produced; in other words, the ideal bacterial cell does not evolve. In the real world, there are cell-cycle process errors that arise from various sources, particularly from unexpectedly variable environments (composition, temperature, irradiation, etc.) and from direct interactions (contact) with other cells.

### 4.2. The process errors of the cell cycle

When the process errors in the cell cycle are *negligibly small*, one cellular micro-reactor will give rise to two copies with vanishingly small differences – only *ideal biological* cells are then produced, and the micro-reactor will not have effectively evolved. In chemical engineering language, the production is within the product’s specifications, for example the level of unreacted starting materials (and other impurities) is acceptably small.

Larger process errors (but non-fatal) make the difference between the two copies quite apparent – the bacterial micro-reactor will have evolved, yielding one or both new cells somewhat dissimilar to the original cell; in chemical engineering language the product is out of specification – it is a different product, which could have better or worse characteristics compared to the original (but in any case it cannot be sold in normal manner!); in bacteriological language, this means that the cellular DNA has changed, producing different strains or phenotypes – the essence of biological evolution (*cf*. Griffith’s experiment with *Streptococcus pneumonia*, indicating a large scale DNA change by natural transformation; or a single nucleotide mutation, A→T in the GAG→GTG codon incorporating glutamic acid instead of valine in the sickle cell hemoglobin).

Very large errors bring about catastrophic failures – the death of bacterial cells; in chemical engineering language, the reactor becomes non-operational (may even cause an industrial accident), as it was not designed to cope with such unexpected (erroneous) inputs. Such ‘fatal’ errors may come from large and sudden changes in the environment when the process control computer’s capability becomes compromised – or in Darwinian language, the genome’s performance has not yet reached the required evolutionary stage to successfully cope with new conditions, *i.e*. to ensure its own survival; the bacterial ‘species’ may then die out if new environmental conditions prevail for too long. It is not inconceivable that large errors may, albeit very rarely, bring about significantly different but still functioning (perhaps poorly) cellular micro-reactors that can ‘dynamically re-stabilize’ and evolve into new kinds of cells, *e.g*. by ‘fusion’ of simple cells, incorporating internal lipid surfaces and multiple chromosomes. Eukaryotic cells did likely arise this way (Sagan, 1967); their multicellular assemblies are now ‘hi-tech’ (highly evolved) engineering designs of ‘wheels-within-wheels’, based on the unit bacterial cell design from a long time ago.

Darwinian evolution then appears naturally as a consequence of non-ideal cellular reproduction – an ‘engineering realization’ of an open, non-equilibrium (non-linear) thermodynamic process, the results of which, for a given genome, are contingent upon physicochemical and biological characteristics of the environment.

### 4.3. Is Darwinian evolution directional toward increasing complexity?

There is little doubt that over the last 3.5 billion years, biological evolution has progressed toward more and more complex cells and multicellular organisms – toward greater complexity, or Darwin’s ‘endless forms most beautiful’, as if by intentional design. The chemical engineering comparison between the error-prone bacterial cell cycle and error-prone chemical manufacturing processes explains qualitatively the apparently intentional direction of evolution. Over the last 3.5 billion years there were many more ‘fatal’ cell cycle errors (there were no ‘intelligent’ chemical engineers) that led to the extinction of many cells and organisms; only a small fraction of minor non-fatal errors (and very, very few *major* non-fatal errors) led to an ‘improved’ physicochemical performance of the cell, or to new kinds of cells; such ‘improvements’ allowed greater chances of maintaining cells’ chemical and physical identity (survival) over an increasing range of extreme environmental gradients (or ‘insults’ in biological language); the cytoplasmic homeostasis has evolved in this way– the maintenance and regulated restructuring of the cytoplasm during variable rates of growth under changing environmental conditions.

Darwinian evolution is thus an *inevitable* result of physicochemical processes, and the direction toward the complex and the sophisticated is *natural*, together with the ‘less appealing’ corollary – the dying-out of less evolvable (physicochemically) cells and organisms, or them ending up as ‘living fossils’ evolutionarily stuck in particular environmental niches. The longer the evolutionary time of ‘fit-for-life’ environments (Henderson, 1913) – now at 3.5 billion years and ticking, the greater the chances of ‘still better’ multicellular systems keeping on evolving.

### 4.4. The future of *Homo sapiens’s* evolution

With the advent of synthetic biology, however, the contingent unintentional Darwinian evolution will cease in the next 1000 years, as *intentionally* designed genetic improvements will become the accelerating evolutionary mechanism for *Homo sapiens* (Church and Regis, 2012). More research is needed on how the brain works as a highly evolved ‘computer *wetware*’, as well as the neuronal connections between the brain and the sensory organs that monitor and respond to the environment. ‘Watson’, the *hardware* solid-state computer, did great playing *Jeopardy* – but will ‘he (or she or it)’ think of something tomorrow? (Bray, 2009; 2012)

Other dramatic variations on the ‘wet’ version of the ‘Watson’ theme are likely to materialize soon, and are in fact currently in progress. For example, the original robots (superficially unrecognizable from humans), their body parts being made in ‘cooking’ vats in the play R.U.R. (*Rossum’s Universal Robots*), are beginning to be synthesized: *cf*. advances in research on human embryonic stem cell (Yirme et al., 2008); another non-Darwinian evolutionary track is being made possible via *Robocops* and similar ‘cyborg’ manifestations: *cf*. current progress in ‘electrical’ interfacing of the brain of the disabled with new mechanical (solid-state) robotic arms. (Hochberg et al. 2012).

## 5. Conclusions and perspectives

Biochemical biotic soups represent realistic (and inexpensive) experimental systems to study the cycling (non-linear, non-equilibrium) biological complexity and the origin of living states through the action of cycling physicochemical gradients. Even though this approach resembles Fred Hoyle’s parable of a rotating tornado trying to assemble a Boeing aircraft from its junked parts (a non-chemical non-sequitur in any case), the experimental goal of assembling a living state from *actual* (not necessarily synthetic) bacterial parts seems worthwhile: to learn how to transit from ‘dead’ to ‘being alive and reproducing’ and *vice versa*. The advantage of the biotic soup as a complex experimental system is that it is ‘close’ to real living systems in its chemical composition. There is a great number of possible designs of ‘biotic soup’ experiments; a combined bacteriological, biochemical, biophysical and biotechnological expertise is needed – particularly in practical aspects of ‘closed membrane, battery-like’ bioenergetics (chemiosmotics) and its interactions with the genetics of the nucleoids, plasmids and phages.

The biotic soup experimentation will motivate the development of new protocols to a) terminate bacterial life to ‘different degrees’, b) to clearly understand the molecular and physicochemical causes of bacterial ‘death’, and c) to identify systems in which ‘living states’ could be detected under cycling non-equilibrium conditions. In principle, we could discover how to ‘stoke’ a multicomponent, multiphase and crowded biochemical system into a dynamic semi-biological existence, perhaps in the form of (fast evolving) progenotes.

## ACKNOWLEDGEMENTS

I thank: Bert Poolman for discussions and collaboration on the structuring of bacterial cells via physicochemical forces, particularly of ionic nature; Elio Schaechter for interest and advising on the relationships between the growth rates of bacteria and their physical (morphological) appearance, and for his help with my blog contribution at http://schaechter.asmblog.org/schaechter/2013/03/feynman-said-just-look-at-the-thing-.html; Dave Deamer for help with the prebiotic aspects of emergent cells and for introducing me to the extensive prebiotic literature; Meir Lahav for his comments on the concept of the cyclic coevolution of biological chirality with the excess of potassium ions in the cytoplasm (Spitzer, 2013); and Gary Pielak and Allen Minton for discussions on biomacromolecular crowding *in vivo*. Wendy Spitzer and Gary Pielak are thanked for help with the manuscript, and MCP Inc. is thanked for support.

